# Biomarkers of Cardiovascular Toxicity of Benzene Inhalation in Mice

**DOI:** 10.1101/2021.08.31.458364

**Authors:** Marina V. Malovichko, Wesley T. Abplanalp, Samantha A. McFall, Breandon S. Taylor, Nalinie S. Wickramasinghe, Israel D. Sithu, Igor N. Zelko, Shizuka Uchida, Saurin R. Sutaria, Michael H. Nantz, Aruni Bhatnagar, Daniel J. Conklin, Timothy E. O’Toole, Sanjay Srivastava

## Abstract

Benzene is a ubiquitous environmental pollutant. Recent population-based studies suggest that benzene exposure is associated with an increased risk for cardiovascular disease. However, it is unclear whether benzene exposure is sufficient to induce cardiovascular toxicity. We examined the effects of benzene inhalation (50 ppm, 6 h/day, 5 days/week, 6 weeks) or HEPA-filtered air exposure on the biomarkers of cardiovascular toxicity in male C57BL/6J mice. Benzene inhalation significantly increased the biomarkers of endothelial activation and injury including endothelial microparticles, activated endothelial microparticles, endothelial progenitor cell microparticles, lung endothelial microparticles, and activated lung and endothelial microparticles while having no effect on circulating levels of endothelial adhesion molecules, endothelial selectins, and biomarkers of angiogenesis. To understand how benzene may induce endothelial injury, we exposed human aortic endothelial cells to benzene metabolites. Of metabolites tested, *trans,trans*-mucondialdehyde (10 μM, 18h) was most toxic. It induced caspases-3, −7 and −9 (intrinsic pathway) activation, and enhanced microparticle formation by 2.4-fold. Levels of plateletleukocyte aggregates, platelet macroparticles, and proportion of CD4^+^ and CD8^+^ T-cells were also significantly elevated in the blood of the benzene-exposed mice. We also found that benzene exposure increased the transcription of genes associated with endothelial cell and platelet activation in the liver; and induced inflammatory genes and suppressed cytochrome P450s in the lungs and the liver. Together, these data suggest that benzene exposure induces endothelial injury, enhances platelet activation and inflammatory processes; and circulatory levels of endothelial cell and platelet-derived microparticles and platelet-leukocyte aggregates are excellent biomarkers of cardiovascular toxicity of benzene.

**Highlights:**

- Inhaled benzene exposure increases the levels of blood endothelial microparticles.
- *In vitro*, benzene metabolite *trans, trans*-mucondialdehyde induces endothelial cell apoptosis and microparticles formation.
- Inhaled benzene exposure decreases the levels of hematopoietic progenitor cells in the bone marrow.
- Inhaled benzene exposure augments the circulating levels of platelet-leukocyte adducts.

## INTRODUCTION

Environmental pollution accounts for 9 million pre-mature deaths worldwide, and two-third of these deaths are attributed to air pollution (1). Benzene, a volatile organic compound (VOC), is abundant both in outdoor and indoor air. Ranked number sixth on the Agency for Toxic Substances and Disease Registry (ATSDR) priority list, benzene is one of the top twenty chemicals generated by industrial sources in the United States. It is used to produce industrial chemicals, rubbers, dyes, lubricants, detergents, etc. (2). The United States Occupational Safety and Health Administration has set the occupational benzene exposure limit of 1 ppm (3), however, benzene exposure in excess of 100 ppm is still prevalent in the developing countries (4). High levels of benzene (>50 ppm) are also generated by tobacco products such as water pipes, cigars, pipe tobacco, and cigarettes (5, 6). Petroleum products and automobile exhaust also contain copious amount of benzene, especially near the emission source (7–9). Indoor sources of benzene include vapor or gases released by benzene containing products such as paints, furniture wax, and detergents (2). The atmospheric benzene exposure is likely to be higher in people living near gasoline refineries, petrochemical industries, gasoline fueling stations, and Superfund and other hazardous waste sites.

Excessive rates of type 2 diabetes and stroke have been found in an evaluation of 720,000 individuals living within a half-mile of 258 Superfund sites that were associated with excessive VOC (such as benzene and trichloroethylene) exposure (10). We observed that environmental benzene exposure is associated with increased CVD risk scores and augmented levels of sub-clinical markers of cardiovascular disease (11–14). Others have shown that benzene exposure increases the risk for arterial hypertension (15, 16), rhythm abnormalities (15), and heart failure(17). An assessment of excessive amount of VOC exposure and cardiovascular disease (CVD) mortality shows that in a single-pollutant model, benzene, propylene, and xylene are all significantly associated with CVD mortality (18). In a cohort study of intra-urban variation in VOCs and mortality, similar associations were found between CVD mortality and exposure to benzene, hexane, and total hydrocarbon (19). However, it is unclear whether benzene exposure is sufficient to cause cardiovascular disease or injury. Therefore, using a well-controlled mouse model, we systematically examined the effect of inhaled benzene exposure on biomarkers of cardiovascular toxicity.

## MATERIALS AND METHODS

### Murine Benzene Exposure

Seven-week-old male C57BL6/J mice were obtained from Jackson laboratories, Bar Harbor, ME. Mice were treated according to American Physiological Society *Guiding Principles in the Care and Use of Animals*, and all protocols were approved by University of Louisville Institutional Animal Care and Use Committee. Mice (n=24/group) were housed under pathogen-free conditions in the University of Louisville vivarium under controlled temperature and 12h light/12h dark cycle. Mice were maintained on a standard chow diet (Rodent Diet 5010, LabDiet, St. Louis, MO) containing 4.5% fat by weight). Starting at eight weeks of age mice were exposed to 50 ppm benzene (6 h/day, 5days/week) for 6 weeks as described before(20, 21). Mice exposed to HEPA-filtered air only served as a control. To examine the effect of benzene exposure on the susceptibility to inflammation, a sub-set of benzene and air-exposed mice (n=24/group) were treated with 0.5 mg/kg lipopolysaccharides (LPS, Sigma Cat# 2630, Lot# 028M4022V; i.p). At the end of the exposure protocol, mice were euthanized with sodium pentobarbital (150 mg/kg body weight; i.p.) and blood and tissues were harvested.

### RNA-seq analysis

One μg of DNase-I-treated total RNA, isolated from the liver and lung tissues, was used for the cDNA library construction for poly-A RNA-seq at Novogene, Sacramento, CA using NEBNext Ultra II RNA Library Prep Kit for Illumina (New England BioLabs, #E7775) according to manufacturer’s protocol. After a series of terminal repair, poly-adenylation, and sequencing adaptor ligation, the double-stranded cDNA library was completed following size selection and PCR enrichment. The resulting 250-350 bp insert libraries were quantified using a Qubit 2.0 fluorometer (Thermo Fisher Scientific) and quantitative PCR. The size distribution was analyzed using an Agilent 2100 Bioanalyzer. Qualified libraries were sequenced on an Illumina Novaseq 6000 system using a paired-end 150 run (2×150 bases). A minimum of 20 million raw reads were generated from each library. The RNA-seq data generated in this study were deposited in the Gene Expression Omnibus (GSE).

For data analysis, FASTQ files were trimmed using fastp(22) (version 0.20.0) and the following parameters: --cut_by_quality5 --cut_by_quality3 --detect_adapter_for_pe --overrepresentation_analysis --correction --trim_front1 7 --trim_front2 7. The alignment to the mouse genome (GRCm38.90) was performed using STAR(23) (version 020201) and the following parameters: --runMode alignReads --runThreadN 8 --outSAMstrandField intronMotif -- outSAMmode Full --outSAMattributes All --outSAMattrRGline --outSAMtype BAM SortedByCoordinate --limitBAMsortRAM 45000000000 --quantMode GeneCounts -- outReadsUnmapped Fastx -outSAMunmapped within. Differential expression analysis was performed using edgeR(24) (version 3.10). The RLE (relative log expression) method was used to normalize the data. Gene ontology analysis was performed with the DAVID Bioinformatics Resources 6.8 (25). A Venn diagram was drawn using Bioinformatics & Evolutionary Genomics website (26). Heatmaps were created with Multiple Experiment Viewer (MEV) (27).

### Bone marrow derived stem cells

Bone marrow cells were isolated from the femur and tibia and separated by Ficoll gradient. The cells were washed twice with PBS containing 1% BSA (PBS/BSA) and incubated with Fc Block (CD32/CD16) anti-mouse antibody for 10 minutes at 4°C to prevent non-specific binding. Samples were incubated (30 min, 4°C) with antibody cocktails containing Lineage-Pacific Blue, ckit-APC-Cy7, Sca-FITC, and CD34-Alexa Fluor 700 antibodies, and analyzed on an LSR II flow cytometer for 90 seconds on high speed. Cell populations were gated using the FlowJo software and normalized to the total number of cells.

### Microparticles

Microparticles in the peripheral blood were measured as described before (28, 29) with slight modifications. Briefly, plasma was centrifuged for 2 min (11,000*xg* at 4°C) to remove residual cells and debris, and the supernatant was aspirated and centrifuged for 45 min (17,000*xg* at 4°C). The resulting microparticle pellet was resuspended in Annexin V Buffer pre-filtered through 0.22μm syringe filter and incubated with the anti-mouse FcBlock (CD32/CD16) for 10 minutes. Endothelial microparticles were stained with the antibody cocktail containing Annexin V-Pacific Blue, Flk-APC, Sca-PECy7, CD62E-PE and CD143-FITC for 30 min. Platelet microparticles were stained in a separate tube with Annexin V-Pacific Blue and platelet CD41-FITC antibody. Identical samples with no antibodies were utilized as controls for the gating. Counting beads, added to individual samples were used for data normalization. Samples were analyzed on BD LSR II flow cytometer for 5 min at low speed. Microparticle numbers were quantified in gated populations <1μm in size and positive for Annexin V staining using the FlowJo software. Microparticle subpopulations were further identified based on expression of various surface markers.

To examine the effect of benzene metabolites on endothelial cell apoptosis and microparticles formation *in vitro*, human aortic endothelial cells (HAEC) were incubated with hydroquinone - HQ, Catechol - Cat, and MA (10 μM each) for 18h, and the apoptosis was examined by western blotting using anti-cleaved-caspase-3, -cleaved caspase-7, -cleaved caspase-8, and cleaved caspase-9 antibody (Cell Signaling Technology, Danvers, MA). The microparticles (<1μm, Annexin V^+^) released in the cell culture medium were analyzed by flow cytometry.

### Synthesis of *trans,trans*-Mucondialdehyde

*trans,trans*-Mucondialdehyde (MA) was prepared from muconic acid (Sigma-Aldrich, St. Louis, MO) by a recently developed one-pot acid-to-aldehyde reduction protocol (30)

### Markers of endothelial function and inflammation

Levels of soluble adhesion molecules, markers of angiogenesis, and cytokines and chemokine in the plasma were measured by multiplex arrays at Eve Technologies (Calgary, Alberta, Canada).

### Immune cells

Circulating immune cells were analyzed by flow cytometry as described before (28, 31). Briefly, lysed whole blood was centrifuged and washed twice with PBS containing 1% BSA (PBS/BSA). The cell pellets were re-suspended in the same buffer and incubated with CD32/CD16 for 10 min at 4°C to prevent unspecific binding. The cells were then incubated with an antibody cocktail consisting of FITC-anti-Nk1.1, PE-anti-Ly6C, PerCPe710-anti-CD8, PECy7-anti-CD62, APC-anti-CD19, Alexa 700-antiGr-1, APCe780-anti-CD3, eVolve605-CD11b, and e650-anti-CD4. After 30 min on ice, the cells were washed, re-suspended in PBS/BSA, and analyzed on an LSR II flow cytometer for 90 sec on high speed. Cell numbers were analyzed using the FlowJo software and normalized to the total leukocyte numbers.

### Platelet-leukocyte aggregates

Platelet-leukocyte aggregates were identified by flow cytometry and quantified as events double positive for CD41 (platelets) and CD 45 (leukocytes) as described before(28).

### Statistics

Data are expressed as mean ± standard error of mean (SEM). Statistical significance was accepted at P<0.05 level. Student’s two-tailed *t* test with unequal variance was used to compare the data sets.

## RESULTS

### Inhaled benzene exposure and endothelial microparticles formation

The endothelium is a critical regulator of vascular homeostasis, vascular tone, angiogenesis, and thrombosis. Our recent studies demonstrate that exposure to benzene depletes circulating endothelial progenitor cells (EPCs, also known as circulating angiogenic cells) in human and mice (11, 14). Mobilization of EPCs from the bone marrow and their homing to the injury sites can be affected by exogenic factors such as aging, disease, and an unhealthy lifestyle (32–36), and therefore changes in EPC levels are reflective of endothelial health. A decrease in the levels of blood EPCs reflects endothelial injury and impaired repair. To assess the benzene exposure-induced endothelial toxicity, we measured circulating endothelial microparticles. Endothelial microparticles, 0.1-1.0 μm vesicles shed from activated or injured cells, are surrogate markers of endothelial activation and injury and comprise 5-15% of microparticles in the blood. Circulating endothelial microparticles are positively associated with coronary artery disease and stroke (37, 38). We observed that inhaled benzene exposure significantly increases the levels of circulating endothelial microparticles (<1μm; Annexin V^+^/CD41^-^/Flk^+^), activated endothelial microparticles (<1μm; Annexin V^+^/CD41^-^/CD62E^+^ [E-selectin]), EPC-derived microparticles (<1μm; Annexin V^+^/CD41^-^/Flk^+^/Sca^+^), lung endothelial microparticles (<1μm; Annexin V^+^/CD143^+^), and activated lung endothelial microparticles (<1μm; Annexin V^+^/CD143^+^/CD62E^+^) by 1.8-3.8-fold (**Fig. 1**). To assess the effect of inhaled benzene exposure on endothelial activation we measured the circulating levels of soluble adhesion molecules. Our data showed that benzene exposure modestly decreased the levels of soluble intra cellular adhesion molecule-1 (sICAM-1; ***Supplemental Table 1***), whereas other soluble adhesion molecules - platelet endothelial cell adhesion molecule-1 (sPECAM-1), endothelial selectin (sE-selectin), and platelet selectin (sP-selectin) were comparable with air-exposed controls. Together these data suggest that inhaled benzene exposure does not induce endothelial activation and most of the endothelial microparticles are derived from benzene-induced endothelial injury. Since benzene inhalation also depletes circulating angiogenic cells mice (11, 14), we also quantified circulating angiogenesis markers. However, our data show that levels of angiogenesis markers in benzene-exposed mice are comparable to the air-exposed controls (***Supplemental Table 1***).

**Figure 1:**
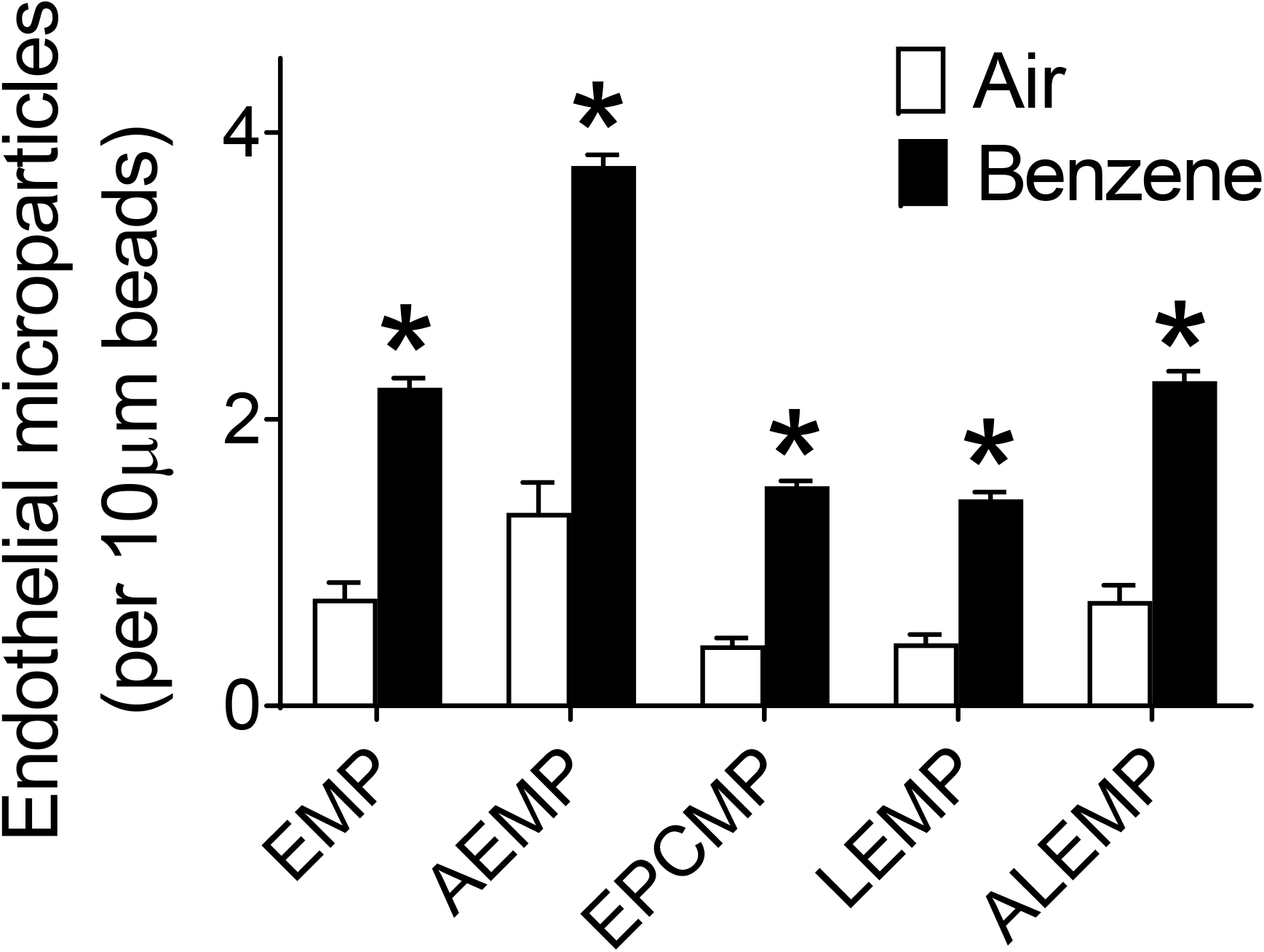
Benzene exposure increases circulating endothelial microparticles in mice. Abundance of endothelial microparticles (EMP; <1μm, AnnexinV^+^/Flk^+^), activated endothelial microparticles (AEMP; <1μm, AnnexinV^+^/CD62E^+^), endothelial progenitor cell microparticles (EPCMP; <1μm, AnnexinV^+^/Flk^+^/Sca^+^), lung endothelial microparticls (LEMP; <1μm, AnnexinV^+^/Flk^+^/CD143^+^), and activated lung endothelial microparticles (ALEMP; <1μm, AnnexinV^+^/CD62E^+^/CD143^+^) in the plasma of benzene- or HEPA-filtered air-exposed mice were analyzed by flow cytometry (n=10/ group). Values are mean ± SEM. *P<0.05 vs control mice.

Because toxicity of benzene is mediated by its metabolism to reactive metabolites, next we directly examined the effect of benzene metabolites hydroquinone, catechol, and MA on HAEC apoptosis. As shown in **Fig. 2**, hydroquinone and catechol only modestly increased the activation of the pan apoptosis marker caspase-7 in HAEC, whereas MA profoundly increased caspase-7 cleavage. MA also robustly increased caspase-3 activation, suggesting that it is the most toxic benzene metabolite for endothelial cells. To examine the mechanisms by which MA exerts its toxicity, we measured the activation of caspase-8 and caspase-9. As shown in **Fig. 2**, MA had no effect on caspase-8, but robustly activated caspase-9. Together, these data suggest that MA doesn’t affect the extrinsic pathway of apoptosis, but selectively activates the intrinsic pathway.

**Figure 2:**
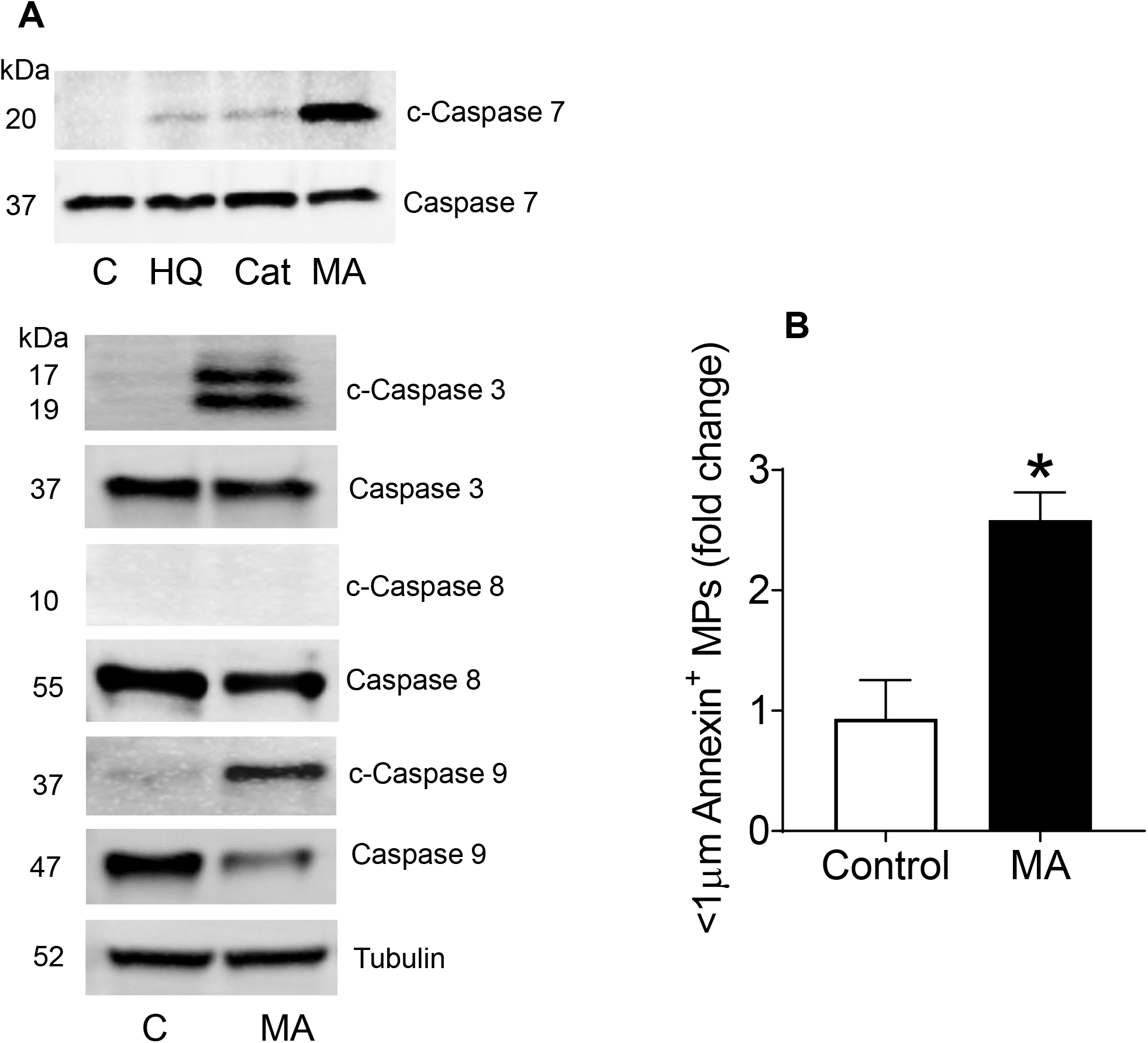
*Benzene metabolite trans,trans*-mucondialdehyde (MA) *increases endothelial microparticle formation from human aortic endothelial cells*. **A**. Caspase activation in human aortic endothelial cells (HAEC) incubated with benzene metabolites (hydroquinone - HQ, Catechol - Cat, and *t,t-mucondialdehyde* - MA; 10 μM each, 18h). **B.** MA (10 μM, 18h, n=6/group) - induced microparticle formation from HAEC. Values are mean ± SEM. *P<0.05 vs controls.

### Inhaled benzene exposure and hematopoietic progenitor cells

Benzene is a well-known hematopoietic toxin (39). Metabolites of benzene such as hydroquinone, catechol, and MA can diffuse from their sites of generation and exert the toxicity at distal sites. We observed that under our experimental conditions, benzene exposure did not affect the levels of common myeloid progenitor (CMPC) and multipotent progenitor cells (MPC) in the bone marrow but significantly decreased the levels of hematopoietic progenitor cells (HPC; **Fig. 3**). Because the vascular niche of HPC is critical for hematopoiesis and endothelial cell materialization, benzene-inhalation-induced depletion of HPC could affect EPC formation in the bone marrow and compromise endothelial repair.

**Figure 3:**
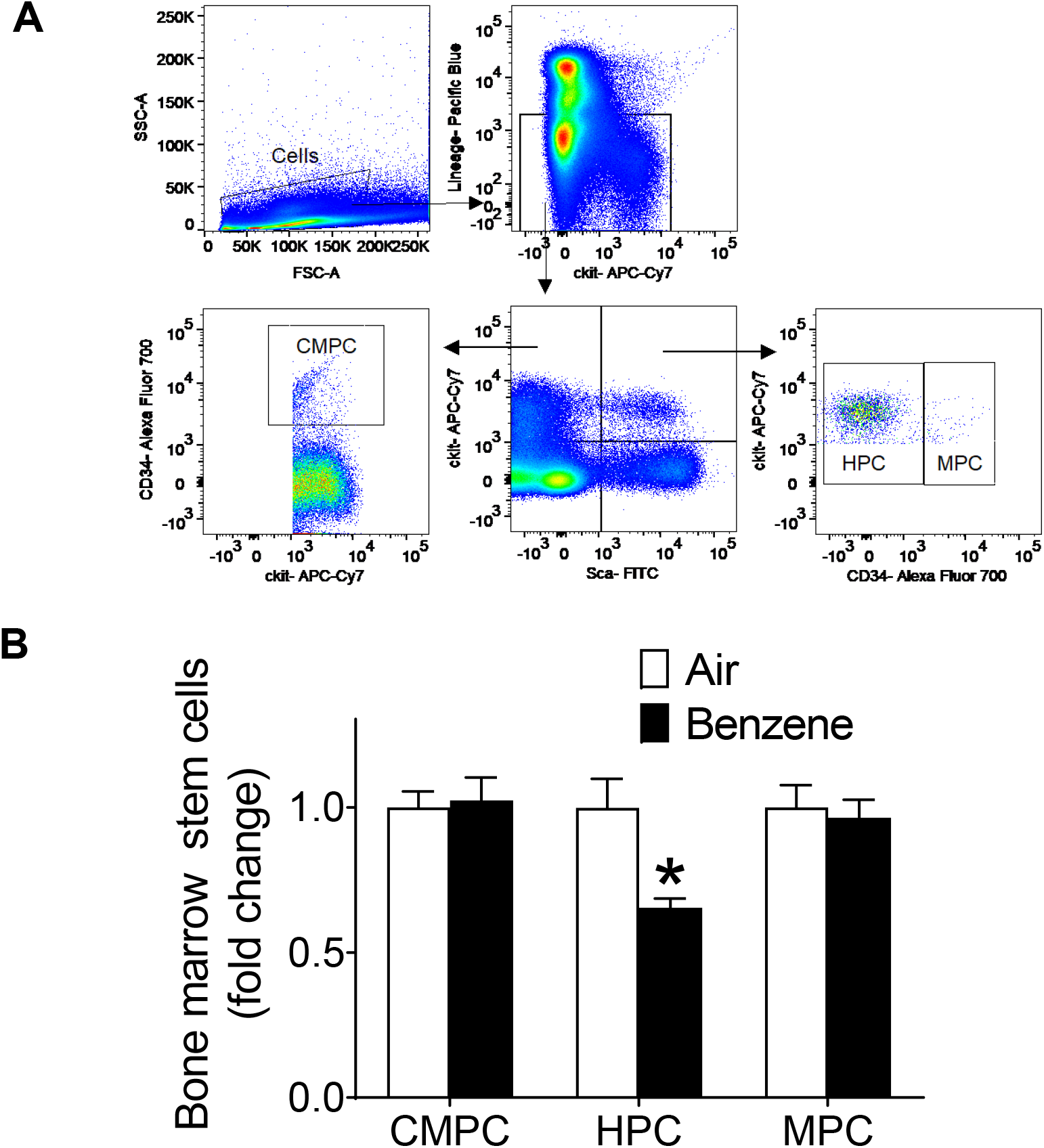
Benzene exposure depletes hematopoietic progenitor cells in the bone marrow. Mice were exposed to benzene or HEPA-filtered air as described under *Methods* and the bone marrow derived stem cells were analyzed by flow cytometry (n=10/ group). Subpopulations of stem cells were identified based on expression of surface markers: Common Myeloid Progenitor Cells (CMPC; Lin^-^ckit^+^Sca^-^CD34^+^), Hematopoietic Progenitor Cells (HPC; Lin^-^ckit^+^Sca^+^CD34^-^) and Multipotent Progenitor Cells (MPC; Lin^-^ckit^+^Sca^+^CD34^+^). Panel **A** depicts the gating scheme for measuring hematopoietic stem cells. Panel **B** shows effects of benzene exposure on bone marrow stem cells. Values are mean ± SEM. *P<0.05 vs control mice.

### Inhaled benzene exposure and platelet activation

The surface of quiescent endothelial cells (luminal surface) which face the blood is normally anti-adhesive. However, injury to the endothelial cells promotes platelet adhesion for repair. We observed that inhaled benzene exposure augments platelet-leukocyte aggregate formation by 3-fold (**Fig. 4**). This was accompanied by 1.6-fold increase in circulating levels of platelet microparticles in benzene-exposed mice. Together these data suggest that benzene exposure enhances platelet activation and platelet-derived microparticles could serve as a biomarker of pro-thrombotic response of inhaled benzene.

**Figure 4:**
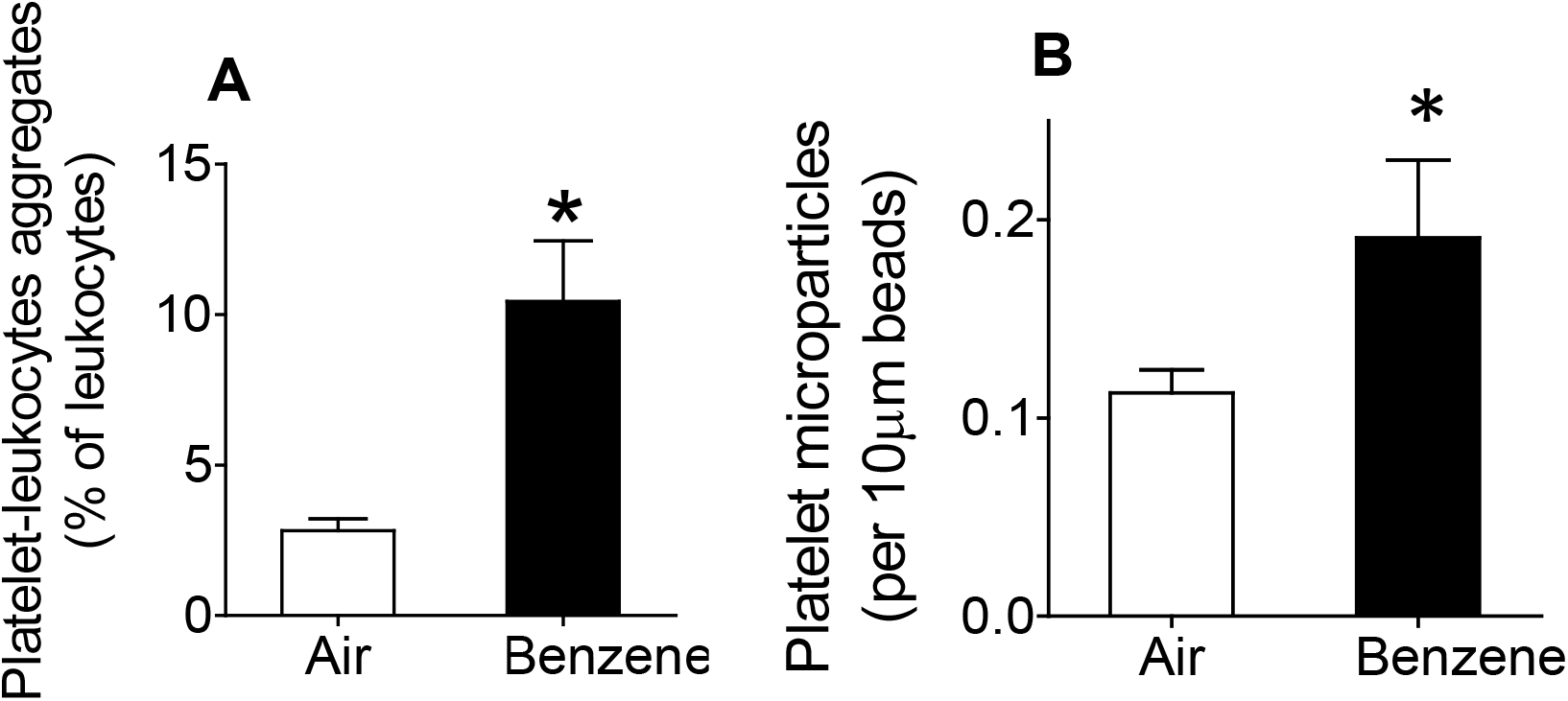
Benzene exposure augments platelet-leukocyte adduct formation. Markers of platelet-leukocyte aggregates were analyzed in the peripheral blood of HEPA-filtered air and benzene-exposed mice by flow cytometry as described under *Methods*. **A**. Platelet-leukocyte adduct (n=10/ group) formation assayed using FITC-labeled anti-CD-41(platelets) and APC-labeled anti-CD 45 (leukocyte) antibodies. **B**. Platelet microparticle levels (< 1 μm cells double positive for Annexin V and CD41). Values are mean ± SEM. *P<0.05 vs control mice.

### Inhaled benzene exposure and inflammatory markers

In humans, polymorphism of cytokines and endothelial activation markers increases the susceptibility to benzene-induced hematopoietic toxicity (40). We have recently shown that in addition to depleting circulating EPCs, benzene inhalation also suppresses the levels of leukocytes, lymphocytes, monocytes, and neutrophils in the peripheral blood in mice (11). However, our flow cytometric analysis of T-lymphocytes shows that inhaled benzene exposure modestly increases the circulating levels of CD3^+^, CD4^+^ and CD8^+^ T-cells (**Fig 5**). Blood CD19^+^ B cells, NK1.1^+^ natural killer cells, Gr1^+^ granulocytes and Ly6C^+^ monocytes (**Fig. 5**) in benzene-exposed mice were comparable to the corresponding air-exposed controls. Quantitation of plasma cytokines showed that IL-6 levels were significantly lower in benzene-exposed mice. All the other circulating cytokines in the benzene-exposed mice were comparable with the air-exposed controls (***Supplemental Table 1***). Stimulation with low dose LPS, 18h before euthanasia, significantly increased the levels of cytokines and chemokines such as GM-CSF, IL-6, IL-10, KC, MCP-1 etc. in the peripheral blood. However, benzene exposure did not affect the LPS-induced cytokine formation.

**Figure 5:**
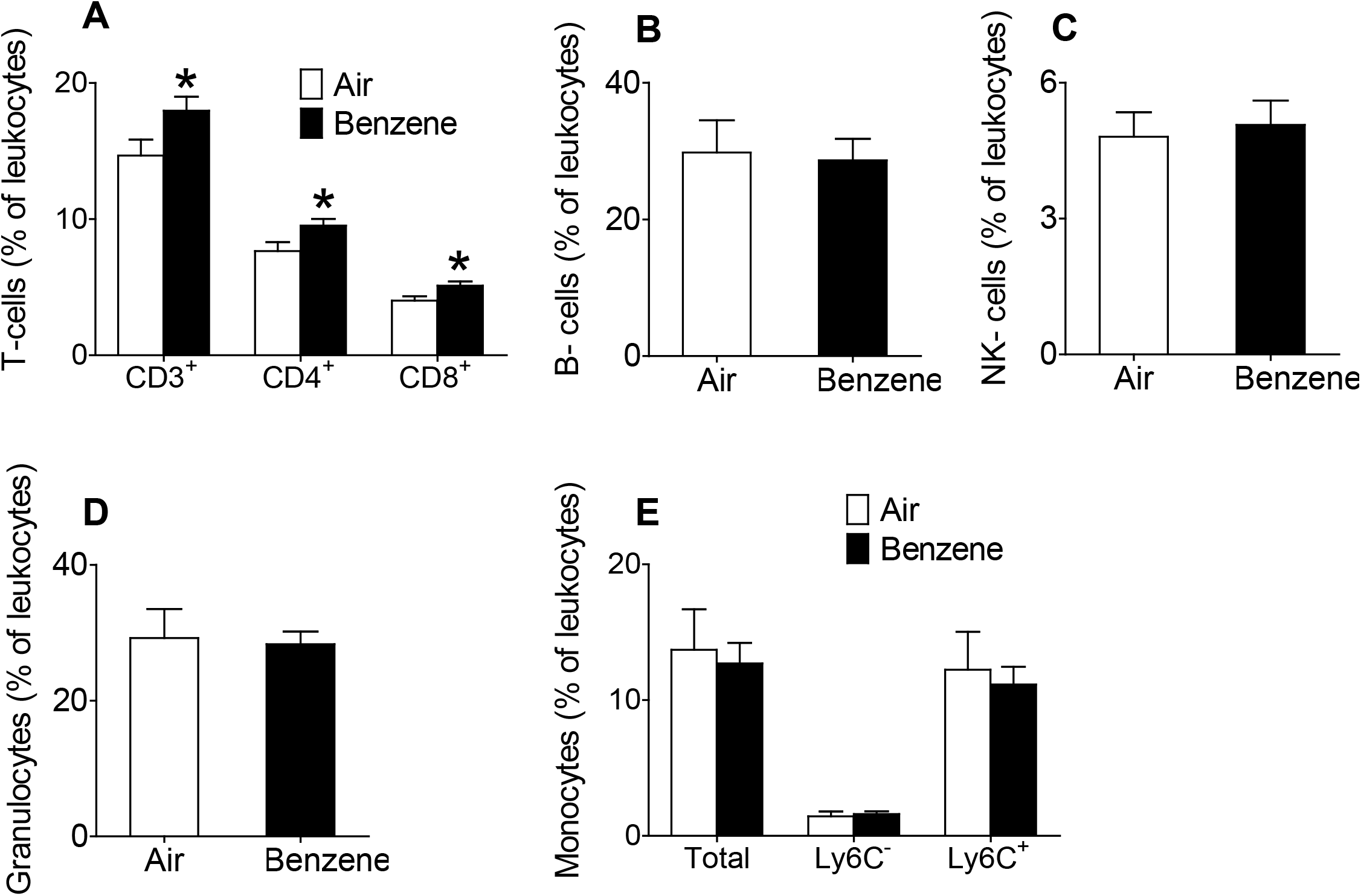
Benzene exposure enhances circulating lymphocytes. Levels of lymphocytes were measured in the peripheral blood by flow cytometry as described under *Methods*. **A**. T-cells (CD3^+^, CD4^+^, and CD8^+^), **B**. B-cells (CD19^+^). **C**. natural killer (NK)-cells (NK1.1^+^). **D**. Granulocytes (GR1^+^). **E**. Monocytes (CD11b^+^). Ly6C^-^ and Ly6C^+^ subpopulations were measured by flow cytometry (n=10/ group). Values are mean ± SEM. *P<0.05 vs control mice.

### Inhaled benzene exposure and pulmonary and hepatic metabolism

While benzene is primarily metabolized in the liver, lungs are the first target of inhaled benzene. We, therefore, examined gene transcription in the liver and the lungs of benzene exposed mice. Although six weeks of benzene exposure did not affect the expression of CYP2E1, which plays a pivotal role in benzene metabolism, benzene exposure significantly down-regulated 185 genes in the lungs and 29 genes in the liver, whereas transcription of 301 genes was increased in the lungs and 43 genes in the liver (>1.5-fold and P<0.05, **Fig 6**). The Heat map of genes associated with cardiometabolic toxicity showed the suppression of cytochrome P450s and up-regulation of inflammatory genes in the lungs and the liver, increased transcription of glycolysis associated genes in the lungs, and induction of the transcription of genes associated with oxidative stress, endothelial activation, and platelet activation in the liver (**Fig. 6**). Gene ontology analysis showed strong association with the lipid metabolic process, cardiac contractility genes, and keratinization in the lungs, and activation of NF-kB, inflammatory response, leukocyte cell-cell adhesion and apoptosis in the liver (**Fig. 6**). These observations are consistent with benzene-induced endothelial apoptosis and our recent studies demonstrating that the benzene-induced insulin resistance is mediated by NF-kB activation, inflammatory signaling, and oxidative stress in the liver (12).

**Figure 6:**
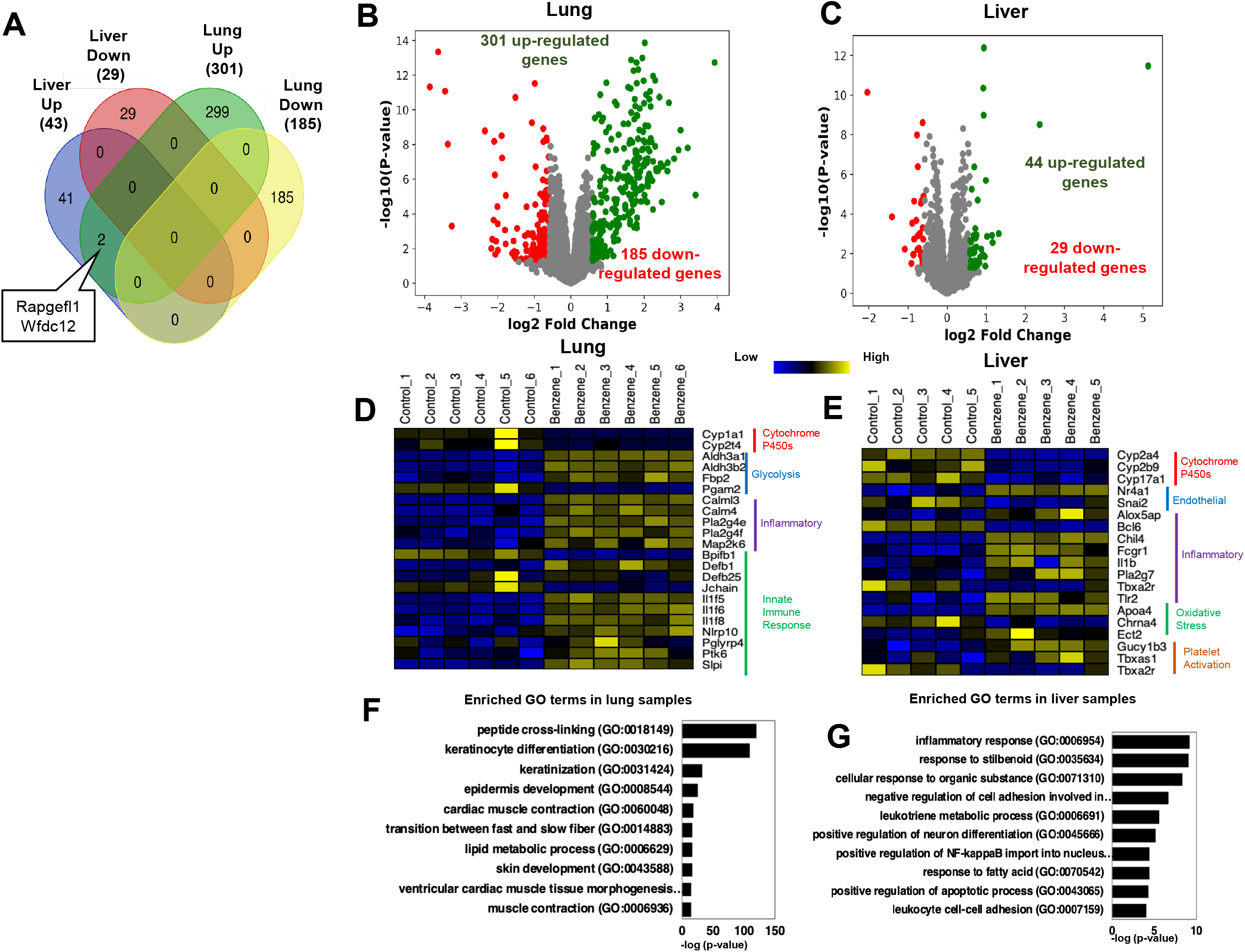
Benzene-induces differential gene regulation in the lung and the liver. Mice were exposed to benzene or HEPA-filtered air as described under *Methods* and RNA-seq analysis was performed on lung and liver tissues (n=6/group). Panel **A** shows the differential regulation of genes in the liver and the lungs of benzene exposed mice. Panels **B** and **C** show the volcano plot of the differentially expressed genes, Panels **D** and **E** illustrate the heat map of prominent gene changes, and panels **F** and **G** depict the gene ontology (GO) analysis of differentially regulated rnRNA in the lungs and the liver, respectively, of benzene-exposed mice.

## DISCUSSION

The major findings of this study are that benzene exposure induces endothelial injury and augments platelet activation, as assessed by a panel of blood endothelial cell and platelet microparticles and platelet-leukocyte adduct formation. This was accompanied by the differential regulation of genes associated with xenobiotic metabolism, endothelial function, platelet activation, and inflammatory signaling associated genes in the liver and the lung, suppression of hematopoietic progenitor cells in the bone marrow, and increase in the levels of T-cells in the peripheral blood.

Although little is known about the direct effect of VOCs such as benzene on vascular injury and thrombosis, the endothelium has been shown to be particularly vulnerable to the effects of tobacco smoke which contains high levels of benzene and other VOCs. In smokers, endothelial dysfunction is the most primitive sign of injury and precedes morphological changes in the vessel wall (41). A dysfunctional endothelium affects vascular homeostasis, blood pressure regulation, thrombosis, atherogenesis, plaque stability, and cardiac functions (42, 43). To examine the effect of benzene exposure on endothelial changes, we measured the levels of microparticles that are released from activated or apoptotic endothelial cells (44) and are a sensitive index of vascular injury (45, 46). Increased levels of circulating endothelial microparticles correlate with endothelial dysfunction in patients with coronary artery disease (36), end stage renal failure (47), obesity (48), and type-2 diabetes (37). Augmented activated endothelial microparticles in the blood are associated with cardiovascular events (49), and enhanced plasma lung endothelial microparticle levels in healthy smokers precede changes in pulmonary function (50). Our data demonstrating that benzene exposure increases the circulating levels of endothelial microparticles, activated endothelial microparticles, EPC microparticles, lung endothelial microparticles and activated lung endothelial microparticles, suggest that these microparticles are sensitive and robust surrogate markers of benzene-induced endothelial injury.

Increased circulating endothelial microparticles have also been observed in humans following episodic fine particulate matter exposure (29). Nonetheless, unlike murine exposure to benzene, fine particulate matter exposure in humans did not increase blood activated endothelial microparticles, suggesting that activated endothelial cells are more sensitive to benzene exposure than fine particulate matter.

Endothelial microparticles contain Von Willebrand factor and factor VIII which promote platelet activation (51, 52). Therefore, the observed increase in platelet-leukocyte adduct formation in benzene-exposed mice could be secondary to benzene-induced endothelial injury and microparticle formation. Moreover, hypercholesterolemia following benzene exposure (11) could also augment platelet activation. Induction of thromboxane A synthase 1 (Tbxas1) in the liver of benzene-exposed mice also corroborates hyper platelet activation, whereas hepatic induction of guanylate cyclase soluble subunit β-1 (Gucyb1), the receptor of nitric oxide, could reflect an adaptive response to benzene exposure-induced platelet activation.

MA-induced endothelial cell apoptosis and endothelial microparticle formation suggest that the observed toxicity of benzene is likely to be mediated by its reactive metabolites. Hepatic induction of the orphan nuclear receptor Nr4a1 (Nurr77), a molecular regulator of apoptosis and inflammation (53–59), in benzene-exposed mice further support that benzene exposure affects apoptotic and inflammatory processes. Although the contribution of benzene-induced Nr4a1 transcription in endothelial toxicity is unknown, Nr4a1 has been suggested to prevent TNFα and IL-1β-induced endothelial activation (60), and endothelial deficiency of Nr4a1suppresses oxLDL-induced apoptosis (57). Unlike Nr4a1, benzene exposure suppressed the hepatic expression of Snai2, a transcription factor involved in the endothelial to mesenchymal transition and implicated in pathological angiogenesis and atherosclerosis (61, 62). Further studies are required to examine the contribution of Snai2 in benzene-induced endothelial toxicity.

Benzene-induced increase in the expression of Thromboxane A synthase 1 (Tbxas1) in the liver corroborate benzene-induced platelet activation. Because thromboxanes play a critical role in modulating vasoconstriction and platelet aggregation, increased formation of thromboxanes can disrupt vascular homeostasis and promote thrombotic vascular events. Increased hepatic transcription of guanylate cyclase 1 soluble subunit beta 1(Gucy1b1) could and guanylate cyclase 1 soluble subunit beta 1(Gucy1b1), the receptor for nitric oxide, could be an adaptive response to mitigate benzene-induced pro-thrombotic responses. However, additional studies are required to examine which cells in the liver induce Tbxas1 and Gucy1b1 and how do these proteins affect benzene-induced vascular homeostasis and thrombosis. Likewise, additional studies are also required to examine the contribution of benzene inhalation-induced transcription of an array of inflammatory genes in the lungs and the liver on endothelial toxicity and platelet activation.

Together, these studies suggest that inhaled benzene exposure induces endothelial injury and affects platelet activation and inflammatory processes. Because benzene is a pervasive and abundant air pollutant, decreasing its exposure can significantly reduce air pollution-induced cardiovascular disease.

---

**Supplemental table1:** Plasma parameters in benzene-exposed mice.

| Parameters (pg/mL) | Air | Benzene | Air + LPS | Benzene + LPS |
| --- | --- | --- | --- | --- |
| **Adhesion molecules** |  |  |  |  |
| sE-Selectin | 643±59 | 610±66 | 2154±161 | 2131±204 |
| sP-Selectin | 1732±254 | 1390±119 | 2427±336 | 2217±328 |
| sICAM-1 | 128±6 | 109±6* | 476±25 | 466±23 |
| Pecam-1 | 37±3 | 30±2 | 48±3 | 41±3 |
| **Angiogenesis markers** |  |  |  |  |
| Angiopoietin-2 | 4732±342 | 4872±243 |  |  |
| EGF | 156±89 | 513±339 |  |  |
| FGF-2 | ND | ND |  |  |
| HGF | 327±74 | 232±26 |  |  |
| Leptin | 608±126 | 801±222 |  |  |
| PLGF-2 | 3.3±0.7 | 3.2±0.4 |  |  |
| SDF-1 | ND | ND |  |  |
| VEGF | 1.02±0.1 | 1.12±0.05 |  |  |
| **Cytokines** |  |  |  |  |
| GM-CSF | 2±1 | 14±8 | 9±2 | 8±1 |
| IFNγ | 1.3±0.4 | 2.3±0.5 | 1.8±0.3 | 1.6±0.4 |
| IL-1α | 8±3 | 3±1 | 14±4 | 6±2 |
| IL-1β | ND | ND | ND | ND |
| IL-2 | 13±6 | 17±11 | 15±8 | 5±1 |
| IL-4 | 0.19±0.11 | 0.04±0.01 | 0.19±0.04 | 0.16±0.04 |
| IL-5 | 6±1 | 5±1 | 13±3 | 9±1 |
| IL-6 | 22±8 | 4±1* | 293±54 | 354±80 |
| IL-7 | 22±9 | 9±5 | 11±5 | 23±9 |
| IL-10 | 3.7±0.4 | 3.4±0.4 | 583.5±39.6 | 447±45.5<sup>#</sup> |
| IL-12 | 6±2 | 6±1 | 17±5 | 11±2 |
| IL-13 | 10±2 | 9±2 | 12±1 | 18±4 |
| LIX | 130±61 | 166±72 | 165±41 | 222±43 |
| IL-17A | 6±2 | 5±1 | 4±1 | 6±1 |
| KC | 406±153 | 178±34 | 3048±443 | 2675±503 |
| MCP-1 | 14±3 | 31±22 | 593±133 | 481±56 |
| MIP-2 | 75±14 | 53±8 | 192±25 | 210±51 |

## References

1. Collaborators GBDRF. Global, regional, and national comparative risk assessment of 79 behavioural, environmental and occupational, and metabolic risks or clusters of risks, 1990-2015: a systematic analysis for the Global Burden of Disease Study 2015. Lancet. 2016;388(10053):1659–724. Epub 2016/10/14. doi: 10.1016/S0140-6736(16)31679-8. PubMed PMID: 27733284; PMCID: PMC5388856.

2. CDC. Toxicological Profile for Benzene. Agency for Toxic Substances and Disease Registry. 2007;CAS#: 71-43-2.

3. Smith MT. Advances in understanding benzene health effects and susceptibility. Annu Rev Public Health. 2010;31:133–48 2 p following 48. doi: 10.1146/annurev.publhealth.012809.103646. PubMed PMID: 20070208; PMCID: PMC4360999.

4. Wong O, Fu H. Exposure to benzene and non-Hodgkin lymphoma, an epidemiologic overview and an ongoing case-control study in Shanghai. Chem Biol Interact. 2005;153-154:33–41. doi: 10.1016/j.cbi.2005.03.008. PubMed PMID: 15935798.

5. Jacob P, 3rd, Abu Raddaha AH, Dempsey D, Havel C, Peng M, Yu L, Benowitz NL. Comparison of nicotine and carcinogen exposure with water pipe and cigarette smoking. Cancer Epidemiol Biomarkers Prev. 2013;22(5):765–72. Epub 2013/03/07. doi: 10.1158/1055-9965.EPI-12-1422. PubMed PMID: 23462922; PMCID: PMC3650103.

6. Appel BR, Guirguis G, Kim IS, Garbin O, Fracchia M, Flessel CP, Kizer KW, Book SA, Warriner TE. Benzene, benzo(a)pyrene, and lead in smoke from tobacco products other than cigarettes. Am J Public Health. 1990;80(5):560–4. Epub 1990/05/01. doi: 10.2105/ajph.80.5.560. PubMed PMID: 2327532; PMCID: PMC1404640.

7. Clayton CA, Pellizzari ED, Whitmore RW, Perritt RL, Quackenboss JJ. National Human Exposure Assessment Survey (NHEXAS): distributions and associations of lead, arsenic and volatile organic compounds in EPA region 5. J Expo Anal Environ Epidemiol. 1999;9(5):381–92. Epub 1999/11/30. doi: 10.1038/sj.jea.7500055. PubMed PMID: 10554141.

8. Morello-Frosch RA, Woodruff TJ, Axelrad DA, Caldwell JC. Air toxics and health risks in California: the public health implications of outdoor concentrations. Risk Anal. 2000;20(2):273–91. Epub 2000/06/22. doi: 10.1111/0272-4332.202026. PubMed PMID: 10859786.

9. Fraser MP, Cass GR, Simoneit BRT. Gas-phase and particle-phase organic compounds emitted from motor vehicle traffic in a Los Angeles roadway tunnel. Environ Sci Technol. 1998;32(14):2051–60. doi: DOI 10.1021/es970916e. PubMed PMID: WOS:000074839400004.

10. Lybarger JA, Lee R, Vogt DP, Perhac RM, Jr., Spengler RF, Brown DR. Medical costs and lost productivity from health conditions at volatile organic compound-contaminated superfund sites. Environ Res. 1998;79(1):9–19. Epub 1998/10/03. doi: 10.1006/enrs.1998.3845. PubMed PMID: 9756676.

11. Abplanalp W, DeJarnett N, Riggs DW, Conklin DJ, McCracken JP, Srivastava S, Xie Z, Rai S, Bhatnagar A, O’Toole TE. Benzene exposure is associated with cardiovascular disease risk. PloS one. 2017; 12(9):e0183602. Epub 2017/09/09. doi: 10.1371/journal.pone.0183602. PubMed PMID: 28886060; PMCID: PMC5590846.

12. Abplanalp WT, Wickramasinghe NS, Sithu SD, Conklin DJ, Xie Z, Bhatnagar A, Srivastava S, O’Toole TE. Benzene Exposure Induces Insulin Resistance in Mice. Toxicol Sci. 2019;167(2):426–37. Epub 2018/10/23. doi: 10.1093/toxsci/kfy252. PubMed PMID: 30346588; PMCID: PMC6358255.

13. Yeager R, Riggs DW, DeJarnett N, Srivastava S, Lorkiewicz P, Xie Z, Krivokhizhina T, Keith RJ, Srivastava S, Browning M, Zafar N, Krishnasamy S, DeFilippis A, Turner J, Rai SN, Bhatnagar A. Association between residential greenness and exposure to volatile organic compounds. Sci Total Environ. 2020;707:135435. Epub 2019/12/23. doi: 10.1016/j.scitotenv.2019.135435. PubMed PMID: 31865083; PMCID: PMC7294698.

14. DeJarnett N, Yeager R, Conklin DJ, Lee J, O’Toole TE, McCracken J, Abplanalp W, Srivastava S, Riggs DW, Hamzeh I, Wagner S, Chugh A, DeFilippis A, Ciszewski T, Wyatt B, Becher C, Higdon D, Ramos KS, Tollerud DJ, Myers JA, Rai SN, Shah J, Zafar N, Krishnasamy SS, Prabhu SD, Bhatnagar A. Residential Proximity to Major Roadways Is Associated With Increased Levels of AC133+ Circulating Angiogenic Cells. Arterioscler Thromb Vasc Biol. 2015;35(11):2468–77. Epub 2015/08/22. doi: 10.1161/ATVBAHA.115.305724. PubMed PMID: 26293462; PMCID: PMC4862408.

15. Kotseva K, Popov T. Study of the cardiovascular effects of occupational exposure to organic solvents. Int Arch Occup Environ Health. 1998;71 Suppl:S87–91. Epub 1998/11/25. PubMed PMID: 9827890.

16. Wiwanitkit V. Benzene exposure and hypertension: an observation. Cardiovasc J Afr. 2007;18(4):264–5. Epub 2007/10/18. PubMed PMID: 17940673.

17. Ran J, Qiu H, Sun S, Yang A, Tian L. Are ambient volatile organic compounds environmental stressors for heart failure? Environ Pollut. 2018;242(Pt B):1810–6. Epub 2018/08/06. doi: 10.1016/j.envpol.2018.07.086. PubMed PMID: 30077408.

18. Tsai DH, Wang JL, Chuang KJ, Chan CC. Traffic-related air pollution and cardiovascular mortality in central Taiwan. Sci Total Environ. 2010;408(8): 1818–23. Epub 2010/02/19. doi: 10.1016/j.scitotenv.2010.01.044. PubMed PMID: 20163830.

19. Villeneuve PJ, Jerrett M, Su J, Burnett RT, Chen H, Brook J, Wheeler AJ, Cakmak S, Goldberg MS. A cohort study of intra-urban variations in volatile organic compounds and mortality, Toronto, Canada. Environ Pollut. 2013. Epub 2013/02/02. doi: 10.1016/j.envpol.2012.12.022. PubMed PMID: 23369806.

20. Blech W. [Alteration of the paper-chromatography determined arginase activity in liver, serum and peritoneal washing fluid in in vivo autolysis of a liver lobe of the rat]. Z Gesamte Exp Med. 1967;144(2):134–44. PubMed PMID: 5590846.

21. Philip A, Tsui A, O’Brien JE. Evaluation of API microtube GT system for detection of germ tube production by clinically significant yeasts. J Clin Microbiol. 1983;18(5):1280–1. doi: 10.1128/JCM.18.5.1280-1281.1983. PubMed PMID: 6358255; PMCID: PMC272886.

22. Chen S, Zhou Y, Chen Y, Gu J. fastp: an ultra-fast all-in-one FASTQ preprocessor. Bioinformatics. 2018;34(17):i884–i90. doi: 10.1093/bioinformatics/bty560. PubMed PMID: 30423086; PMCID: PMC6129281.

23. Dobin A, Davis CA, Schlesinger F, Drenkow J, Zaleski C, Jha S, Batut P, Chaisson M, Gingeras TR. STAR: ultrafast universal RNA-seq aligner. Bioinformatics. 2013;29(1):15–21. doi: 10.1093/bioinformatics/bts635. PubMed PMID: 23104886; PMCID: PMC3530905.

24. Robinson MD, McCarthy DJ, Smyth GK. edgeR: a Bioconductor package for differential expression analysis of digital gene expression data. Bioinformatics. 2010;26(1): 139–40. doi: 10.1093/bioinformatics/btp616. PubMed PMID: 19910308; PMCID: PMC2796818.

25. Huang da W, Sherman BT, Lempicki RA. Systematic and integrative analysis of large gene lists using DAVID bioinformatics resources. Nat Protoc. 2009;4(1):44–57. doi: 10.1038/nprot.2008.211. PubMed PMID: 19131956.

26. Bioinformatics. Calculate and draw custom Venn diagrams. http://bioinformaticspsbugentbe/webtools/Venn/.Bioinformatics & Evolutionary Genomics.

27. Saeed AI, Sharov V, White J, Li J, Liang W, Bhagabati N, Braisted J, Klapa M, Currier T, Thiagarajan M, Sturn A, Snuffin M, Rezantsev A, Popov D, Ryltsov A, Kostukovich E, Borisovsky I, Liu Z, Vinsavich A, Trush V, Quackenbush J. TM4: a free, open-source system for microarray data management and analysis. Biotechniques. 2003;34(2):374–8. doi: 10.2144/03342mt01. PubMed PMID: 12613259.

28. Malovichko MV, Zeller I, Krivokhizhina TV, Xie Z, Lorkiewicz P, Agarwal A, Wickramasinghe N, Sithu SD, Shah J, O’Toole T, Rai SN, Bhatnagar A, Conklin DJ, Srivastava S. Systemic Toxicity of Smokeless Tobacco Products in Mice. Nicotine Tob Res. 2019;21(1):101–10. doi: 10.1093/ntr/ntx230. PubMed PMID: 30085294; PMCID: PMC6329408.

29. Pope CA, 3rd, Bhatnagar A, McCracken JP, Abplanalp W, Conklin DJ, O’Toole T. Exposure to Fine Particulate Air Pollution Is Associated With Endothelial Injury and Systemic Inflammation. Circ Res. 2016;119(11):1204–14. doi: 10.1161/CIRCRESAHA.116.309279. PubMed PMID: 27780829; PMCID: PMC5215745.

30. Sutaria SR, Nantz MH. A Convenient Preparation of Muconaldehyde Using a One-Pot Acid-to-Aldehyde Reduction Protocol. Org Prep Proced Int. 2021;53:In Press.

31. Conklin DJ, Malovichko MV, Zeller I, Das TP, Krivokhizhina TV, Lynch BH, Lorkiewicz P, Agarwal A, Wickramasinghe N, Haberzettl P, Sithu SD, Shah J, O’Toole TE, Rai SN, Bhatnagar A, Srivastava S. Biomarkers of Chronic Acrolein Inhalation Exposure in Mice: Implications for Tobacco Product-Induced Toxicity. Toxicol Sci. 2017;158(2):263–74. doi: 10.1093/toxsci/kfx095. PubMed PMID: 28482051; PMCID: PMC5837482.

32. Heiss C, Amabile N, Lee AC, Real WM, Schick SF, Lao D, Wong ML, Jahn S, Angeli FS, Minasi P, Springer ML, Hammond SK, Glantz SA, Grossman W, Balmes JR, Yeghiazarians Y. Brief secondhand smoke exposure depresses endothelial progenitor cells activity and endothelial function: sustained vascular injury and blunted nitric oxide production. J Am Coll Cardiol. 2008;51(18):1760–71. doi: 10.1016/j.jacc.2008.01.040. PubMed PMID: 18452782.

33. Chang EI, Loh SA, Ceradini DJ, Chang EI, Lin SE, Bastidas N, Aarabi S, Chan DA, Freedman ML, Giaccia AJ, Gurtner GC. Age decreases endothelial progenitor cell recruitment through decreases in hypoxia-inducible factor 1alpha stabilization during ischemia. Circulation. 2007;116(24):2818–29. doi: 10.1161/CIRCULATIONAHA.107.715847. PubMed PMID: 18040029.

34. Rauscher FM, Goldschmidt-Clermont PJ, Davis BH, Wang T, Gregg D, Ramaswami P, Pippen AM, Annex BH, Dong C, Taylor DA. Aging, progenitor cell exhaustion, and atherosclerosis. Circulation. 2003;108(4):457–63. doi: 10.1161/01.CIR.0000082924.75945.48. PubMed PMID: 12860902.

35. Vasa M, Fichtlscherer S, Aicher A, Adler K, Urbich C, Martin H, Zeiher AM, Dimmeler S. Number and migratory activity of circulating endothelial progenitor cells inversely correlate with risk factors for coronary artery disease. Circ Res. 2001;89(1):E1–7. PubMed PMID: 11440984.

36. Werner N, Wassmann S, Ahlers P, Kosiol S, Nickenig G. Circulating CD31+/annexin V+ apoptotic microparticles correlate with coronary endothelial function in patients with coronary artery disease. Arterioscler Thromb Vasc Biol. 2006;26(1):112–6.doi: 10.1161/01.ATV.0000191634.13057.15. PubMed PMID: 16239600.

37. Koga H, Sugiyama S, Kugiyama K, Watanabe K, Fukushima H, Tanaka T, Sakamoto T, Yoshimura M, Jinnouchi H, Ogawa H. Elevated levels of VE-cadherin-positive endothelial microparticles in patients with type 2 diabetes mellitus and coronary artery disease. J Am Coll Cardiol. 2005;45(10):1622–30. Epub 2005/05/17. doi: 10.1016/j.jacc.2005.02.047. PubMed PMID: 15893178.

38. Simak J, Gelderman MP, Yu H, Wright V, Baird AE. Circulating endothelial microparticles in acute ischemic stroke: a link to severity, lesion volume and outcome. J Thromb Haemost. 2006;4(6):1296–302. Epub 2006/05/19. doi: 10.1111/j.1538-7836.2006.01911.x. PubMed PMID: 16706974.

39. McHale CM, Zhang L, Smith MT. Current understanding of the mechanism of benzene-induced leukemia in humans: implications for risk assessment. Carcinogenesis. 2012;33(2):240–52. Epub 2011/12/15. doi: 10.1093/carcin/bgr297. PubMed PMID: 22166497; PMCID: PMC3271273.

40. Lan Q, Zhang L, Shen M, Smith MT, Li G, Vermeulen R, Rappaport SM, Forrest MS, Hayes RB, Linet M, Dosemeci M, Alter BP, Weinberg RS, Yin S, Yeager M, Welch R, Waidyanatha S, Kim S, Chanock S, Rothman N. Polymorphisms in cytokine and cellular adhesion molecule genes and susceptibility to hematotoxicity among workers exposed to benzene. Cancer Res. 2005;65(20):9574–81. doi: 10.1158/0008-5472.CAN-05-1419. PubMed PMID: 16230423.

41. Poredos P, Orehek M, Tratnik E. Smoking is associated with dose-related increase of intima-media thickness and endothelial dysfunction. Angiology. 1999;50(3):201–8. PubMed PMID: 10088799.

42. Lind L, Berglund L, Larsson A, Sundstrom J. Endothelial function in resistance and conduit arteries and 5-year risk of cardiovascular disease. Circulation. 2011;123(14):1545–51. doi: 10.1161/CIRCULATIONAHA.110.984047. PubMed PMID: 21444885.

43. Gokce N. Clinical assessment of endothelial function: ready for prime time? Circ Cardiovasc Imaging. 2011;4(4):348–50. doi: 10.1161/CIRCIMAGING.111.966218. PubMed PMID: 21772010; PMCID: PMC3144162.

44. Mause SF, Weber C. Microparticles: protagonists of a novel communication network for intercellular information exchange. Circ Res. 2010; 107(9): 1047–57. doi: 10.1161/CIRCRESAHA.110.226456. PubMed PMID: 21030722.

45. Philippova M, Suter Y, Toggweiler S, Schoenenberger AW, Joshi MB, Kyriakakis E, Erne P, Resink TJ. T-cadherin is present on endothelial microparticles and is elevated in plasma in early atherosclerosis. Eur Heart J. 2011;32(6):760–71. doi: 10.1093/eurheartj/ehq206. PubMed PMID: 20584775.

46. Pirro M, Schillaci G, Bagaglia F, Menecali C, Paltriccia R, Mannarino MR, Capanni M, Velardi A, Mannarino E. Microparticles derived from endothelial progenitor cells in patients at different cardiovascular risk. Atherosclerosis. 2008;197(2):757–67. doi: 10.1016/j.atherosclerosis.2007.07.012. PubMed PMID: 17720166.

47. Amabile N, Guerin AP, Leroyer A, Mallat Z, Nguyen C, Boddaert J, London GM, Tedgui A, Boulanger CM. Circulating endothelial microparticles are associated with vascular dysfunction in patients with end-stage renal failure. J Am Soc Nephrol. 2005;16(11):3381–8. Epub 2005/09/30. doi: 10.1681/ASN.2005050535. PubMed PMID: 16192427.

48. Esposito K, Ciotola M, Schisano B, Gualdiero R, Sardelli L, Misso L, Giannetti G, Giugliano D. Endothelial microparticles correlate with endothelial dysfunction in obese women. J Clin Endocrinol Metab. 2006;91(9):3676–9. Epub 2006/07/11. doi: 10.1210/jc.2006-0851. PubMed PMID: 16822816.

49. Lee ST, Chu K, Jung KH, Kim JM, Moon HJ, Bahn JJ, Im WS, Sunwoo J, Moon J, Kim M, Lee SK, Roh JK. Circulating CD62E+ microparticles and cardiovascular outcomes. PLoS One. 2012;7(4):e35713. Epub 2012/05/09. doi: 10.1371/journal.pone.0035713. PubMed PMID: 22563392; PMCID: PMC3338519.

50. Gordon C, Gudi K, Krause A, Sackrowitz R, Harvey BG, Strulovici-Barel Y, Mezey JG, Crystal RG. Circulating endothelial microparticles as a measure of early lung destruction in cigarette smokers. Am J Respir Crit Care Med. 2011;184(2):224–32. doi: 10.1164/rccm.201012-2061OC. PubMed PMID: 21471087; PMCID: PMC3172886.

51. Jy W, Jimenez JJ, Mauro LM, Horstman LL, Cheng P, Ann ER, Bidot CJ, Ahn YS. Endothelial microparticles induce formation of platelet aggregates via a von Willebrand factor/ristocetin dependent pathway, rendering them resistant to dissociation. Journal of Thrombosis and Haemostasis. 2005;3(6):1301–8. doi: DOI 10.1111/j.1538-7836.2005.01384.x. PubMed PMID: WOS:000229873700031.

52. Jimenez JJ, Jy W, Mauro LM, Horstman LL, Soderland C, Ahn YS. Endothelial microparticles released in thrombotic thrombocytopenic purpura express von Willebrand factor and markers of endothelial activation. Brit J Haematol. 2003; 123(5):896–902. doi: DOI 10.1046/j.1365-2141.2003.04716.x. PubMed PMID: WOS:000186695700015.

53. Li QX, Ke N, Sundaram R, Wong-Staal F. NR4A1, 2, 3--an orphan nuclear hormone receptor family involved in cell apoptosis and carcinogenesis. Histol Histopathol. 2006;21(5):533–1.Epub 2006/02/24. doi: 10.14670/HH-21.533. PubMed PMID: 16493583.

54. Pei L, Castrillo A, Tontonoz P. Regulation of macrophage inflammatory gene expression by the orphan nuclear receptor Nur77. Mol Endocrinol. 2006;20(4):786–94. Epub 2005/12/13. doi: 10.1210/me.2005-0331. PubMed PMID: 16339277.

55. Hanna RN, Shaked I, Hubbeling HG, Punt JA, Wu R, Herrley E, Zaugg C, Pei H, Geissmann F, Ley K, Hedrick CC. NR4A1 (Nur77) deletion polarizes macrophages toward an inflammatory phenotype and increases atherosclerosis. Circ Res. 2012;110(3):416–27. Epub 2011/12/24. doi: 10.1161/CIRCRESAHA.111.253377. PubMed PMID: 22194622; PMCID: PMC3309661.

56. Zhao Y, Bruemmer D. NR4A orphan nuclear receptors: transcriptional regulators of gene expression in metabolism and vascular biology. Arterioscler Thromb Vasc Biol. 2010;30(8):1535–2.Epub 2010/07/16. doi: 10.1161/ATVBAHA.109.191163. PubMed PMID: 20631354; PMCID: PMC2907171.

57. Li P, Bai Y, Zhao X, Tian T, Tang L, Ru J, An Y, Wang J. NR4A1 contributes to high-fat associated endothelial dysfunction by promoting CaMKII-Parkin-mitophagy pathways. Cell Stress Chaperones. 2018;23(4):749–61. Epub 2018/02/23. doi: 10.1007/s12192-018-0886-1. PubMed PMID: 29470798; PMCID: PMC6045535.

58. Hilgendorf I, Gerhardt LM, Tan TC, Winter C, Holderried TA, Chousterman BG, Iwamoto Y, Liao R, Zirlik A, Scherer-Crosbie M, Hedrick CC, Libby P, Nahrendorf M, Weissleder R, Swirski FK. Ly-6Chigh monocytes depend on Nr4a1 to balance both inflammatory and reparative phases in the infarcted myocardium. Circ Res. 2014;114(10):1611–22. Epub 2014/03/15. doi: 10.1161/CIRCRESAHA.114.303204. PubMed PMID: 24625784; PMCID: PMC4017349.

59. Martinez-Gonzalez J, Badimon L. The NR4A subfamily of nuclear receptors: new early genes regulated by growth factors in vascular cells. Cardiovasc Res. 2005;65(3):609–18. Epub 2005/01/25. doi: 10.1016/j.cardiores.2004.10.002. PubMed PMID: 15664387.

60. You B, Jiang YY, Chen S, Yan G, Sun J. The orphan nuclear receptor Nur77 suppresses endothelial cell activation through induction of IkappaBalpha expression. Circ Res. 2009;104(6):742–9. Epub 2009/02/14. doi: 10.1161/CIRCRESAHA.108.192286. PubMed PMID: 19213954.

61. Hultgren NW, Fang JS, Ziegler ME, Ramirez RN, Phan DTT, Hatch MMS, Welch-Reardon KM, Paniagua AE, Kim LS, Shon NN, Williams DS, Mortazavi A, Hughes CCW. Slug regulates the Dll4-Notch-VEGFR2 axis to control endothelial cell activation and angiogenesis. Nat Commun. 2020;11(1):5400. Epub 2020/10/28. doi: 10.1038/s41467-020-18633-z. PubMed PMID: 33106502; PMCID: PMC7588439.

62. Evrard SM, Lecce L, Michelis KC, Nomura-Kitabayashi A, Pandey G, Purushothaman KR, d’Escamard V, Li JR, Hadri L, Fujitani K, Moreno PR, Benard L, Rimmele P, Cohain A, Mecham B, Randolph GJ, Nabel EG, Hajjar R, Fuster V, Boehm M, Kovacic JC. Endothelial to mesenchymal transition is common in atherosclerotic lesions and is associated with plaque instability. Nat Commun. 2016;7:11853. Epub 2016/06/25. doi: 10.1038/ncomms11853. PubMed PMID: 27340017; PMCID: PMC4931033.

